# ASSOCIATION STUDY OF SIX CANDIDATE GENES WITH MAJOR DEPRESSIVE DISORDER IN THE NORTH-WESTERN POPULATION OF PAKISTAN

**DOI:** 10.1101/2021.03.01.433336

**Authors:** Naqash Alam, Sadiq Ali, Nazia Akbar, Muhammad Ilyas, Habib Ahmed, Arooj Mustafa, Shehzada Khurram, Zeeshan Sajid, Najeeb Ullah, Shumaila Qayyum, Tariq Rahim, Mian Syed Usman, Nawad Ali, Imad Khan, Khola Pervez, BiBi Sumaira, Nasir Ali, Nighat Sultana, Adeel Yunus Tanoli, Madiha Islam

## Abstract

People around the world are currently affected by Major Depressive Disorder (MDD). Despite its many aspects, symptoms, manifestations and impacts, efforts have been made to identify the root causes of the disorder. In particular, genetic studies have concentrated on identifying candidate genes for MDD and exploring associations between these genes and some specific group of individuals. The aim of this research was to find out the association between single nucleotide polymorphisms in 6 candidate genes linked to the neurobiology of major depressive disorder in the North-Western population of Pakistan. We performed a case-control analysis, with 400 MDD and 232 controls. A trained psychiatrist or clinical psychologists evaluated the patients. Six polymorphisms were genotyped and tested for allele and genotype association with MDD. There were no statistical variations between MDD patients and healthy controls for genotypic and allelic distribution of all the polymorphisms observed. Thus, our analysis does not support the major role of these polymorphisms in contributing to MDD susceptibility, although it does not preclude minor impact. The statistically significant correlation between six polymorphisms and major depressive disorder in the studied population was not observed. There are inconsistencies in investigations around the world. Future research, including GWAS and association analysis on larger scale should be addressed for further validation and replication of the present findings.

## Introduction

Major depression is the most prevalent psychiatric disorder; about 450 million people are affected and the predominant cause of the morbidity and disability worldwide [1]. Every year, 6-7% of the population and 16% of people are affected by depression during their lives [2,3]. In a given year, according to the World Health Organization, 9% of women and 5% of men have developed depressive disorder [4]. It is accompanied by the following symptom; sadness, lack of energy and interest, mood and behavior changes and abnormal circadian rhythms. Major depressive disorder is very sever and heterogeneous and can lead to suicide when it not treated [4,5]. In the etiology of major depression, genetic factors clearly play a significant role, as shown by family and twined studies that indicate heritability estimates of 17% to 75% on average at 37% [6].

In the development of psychiatric disorders, both genetic disposition and environmental factors are significant, especially severe psychiatric disorders such as major depression [7]. It is because the cumulative contribution of genetic influence might be stronger for those severe mental disorders [8], and a range of environmental factors seem to result in an increased proportion of cases of serious mental illness [7,9]. The wide “heritability gap” the disparity between heritability of twin studies and heritability of single molecular nucleotide polymorphism (SNP) in serious mental disorders suggests that gene environmental interactions might account for a large proportion of cases [7]. In order to evaluate gene-environment interactions, some researchers have used comparative methods (e.g. adoption studies; twin studies), but these methods are often restricted due to unequal family relationships and because unique environmental factors can interact with unique genetic variants. In this regard it is important to investigate the interactions involving particular molecular genetics variants to understand the causal relationships that develop into severe mental illness [7]. Gene–environment interaction is involved when two distinct genotypes respond to environmental variations in various ways. It can be characterized as the dependence on the genotype of an individual on the impact of an environmental factor, and vice versa [10]. Recent studies have shown specific interactions between genes and the environment in severe psychiatric disorders, including psychosis, TB, MDD, and PTSD. While many environmental factors are thought to play a role in psychiatric disorders, it has been suggested that childhood trauma has a serious impact on brain development which leading permanent functional changes that may enhance the risk of developing mental health problems [11].

## Materials and Methods

### Subjects

A case-control study was conducted to investigate the association of gene polymorphisms with 400 patients and 232 controls with major depressive disorder; the diagnosis of MDD was made based on the Diagnostic and Statistical Manual of Mental Disorders - Fourth Edition (DSM-IV). A psychiatric specialist interviewed the patients. When the patient was diagnosed with MDD, according to DSM-IV guideline, another psychiatric specialist was to confirm the diagnosis. This study included patients who had been diagnosed with MDD by at least two qualified psychiatrists. The study excluded patients with a history of neurological illness or another psychiatric disorder. The 17-item Hamilton Rating Scale for Depression (HAMD-17) was used to assessed the severity of depression. The research included only subjects with a minimum score of 18 on HAMD-17. Patients with serious organic conditions were excluded to prevent cases with secondary depression. The design study was approved by the Ethical Committee of the Hazara University Mansehra, KP Province. Informed consent was obtained from all subjects and their parents. Individual information on occupation, ethnicity, comorbidities, and level of education was recorded. Patient’s history, family history and demographic data of patients and controls are given in Table 1.

**Table 1.** Demographic and clinical characteristics of patients with MDD and healthy controls.

| Variables | Cases |  | Controls |  | Totals |  |
| --- | --- | --- | --- | --- | --- | --- |
|  | n=400 | % | n=232 | % | n=632 | % |
| **Gender** |  |  |  |  |  |  |
| Male | 197 | 48.9 | 121 | 51.7 | 318 | 50.3 |
| Female | 203 | 50.4 | 111 | 47.4 | 314 | 49.68 |
| **Age** |  |  |  |  |  |  |
| 15-30 | 175 | 43.4 | 101 | 43.2 | 276 | 43.7 |
| 31-60 | 189 | 46.9 | 110 | 47.0 | 299 | 47.3 |
| 61 and above | 36 | 8.9 | 21 | 9.0 | 57 | 4.75 |
| **Weight** |  |  |  |  |  |  |
| 30-50 | 38 | 9.4 | 24 | 10.3 | 62 | 9.81 |
| 51-80 | 330 | 81.9 | 193 | 82.5 | 523 | 82.8 |
| 81 and above | 32 | 7.9 | 15 | 6.4 | 30 | 4.75 |
| **Marital Status** |  |  |  |  |  |  |
| Unmarried | 102 | 25.3 | 56 | 23.9 | 158 | 25.0 |
| Married | 280 | 69.5 | 167 | 71.4 | 447 | 70.7 |
| Divorced | 16 | 4.0 | 7 | 3.0 | 23 | 3.64 |
| Widow | 2 | 0.5 | 2 | 0.9 | 4 | 0.63 |
| **Education** |  |  |  |  |  |  |
| Illiterate | 176 | 43.7 | 101 | 43.2 | 277 | 43.8 |
| Primary | 66 | 16.4 | 40 | 17.1 | 106 | 16.8 |
| Grade 10 | 65 | 16.1 | 36 | 15.4 | 101 | 16.0 |
| Grade 12 | 51 | 12.7 | 35 | 15.0 | 86 | 13.6 |
| Undergraduate | 17 | 4.2 | 7 | 3.0 | 24 | 3.80 |
| Post graduate | 25 | 6.2 | 13 | 5.6 | 38 | 6.01 |
| **Occupation** |  |  |  |  |  |  |
| Employed | 50 | 12.4 | 32 | 13.7 | 82 | 13.0 |
| Unemployed | 302 | 74.9 | 174 | 74.4 | 476 | 75.3 |
| Students | 48 | 11.9 | 26 | 11.1 | 74 | 11.7 |

### Selection of SNPs

The candidate SNPs of genes met at least two criteria from the following: (1) The SNPs must have been previously associated with MDD or a depression episode; (2) The SNPs were located in the intronic or exonic region within genes; and (3) A minor allele frequency of *>*5% in north-western population. The public domain of the database for SNPs of the National Center for Biotechnology Information (NCBI dbSNP), accessible at http://www.ncbi.nlm.nih.gov/snp(Bethesda, Montgomery County, MD, USA), was used to select the studied polymorphisms and to be localized either in the coding or regulatory area of the genes (Table 2).

**Table 2.** Characteristics of the studied polymorphisms.

| Gene | SNP ID | Position | Polymorphism | Localization |
| --- | --- | --- | --- | --- |
| *GSK3B* | rs334535 | Chr:3:120073457 | A/G | Intron |
| *TPH1* | rs1800532 | Chr:11:18026269 | A/C | Intron |
| *TPH2* | rs7305115 | Chr:12:7197908 | A/G | Exon |
| *DAT1* | rs40184 | Chr:5:1394962 | A/G | Intron |
| *SLC6A15* | rs1545843 | Chr:12:84170289 | A/G | Intron |
| *BDNF* | rs6265 | Chr:11:27658369 | G/A | Exon |

### DNA extraction and nucleotide sequences

5 ml of whole blood was collected from each participant and placed in the anticoagulant EDTA tubes. Before processing, samples were saved at −20 ° C. The standard phenol chloroform method was then used to extract genome DNA. The polymerase chain reaction (PCR) method was used for DNA amplification. The primers were designed using by Primer 5.0 software for PCR amplification, and their specificity was validated using NCBI BLASTN (http://www.ncbi.nlm.nih.gov/BLAST/). PCR amplification was performed with a reaction volume of 25 µl containing 2.5 µl buffer 109 (Tiangen, Beijing, China), 200 µM each dNTP, µM primers each, and 1.0 unit of Taq DNA polymerase (Tiangen, Beijing, China) and 60 ng of genomic DNA. Conditions used for PCR amplification were denaturation at 95°C for 10 min, followed by 35 cycles of 95°C for 30 s, 54–56°C for 30 s, and 72°C for 30 s, followed by a final extension at 72°C for 12 min. Table 3 contains primer sequences used for amplification of DNA fragments containing polymorphisms. PCR products obtained were commercially sequenced by Macrogen Korea. Sanger sequencing analysis was performed on a comparative basis, where the patient’s electropherogram was compared against an electropherogram from healthy/control samples. The observed differences between the two groups was recorded and analyzed. DNA sequence was analyzed using ClustalW multiple sequence alignment (Bioedit, v7.0.9; available at http://www.mbio.ncsu.edu/BioEdit/page2.html).

**Table 3.**
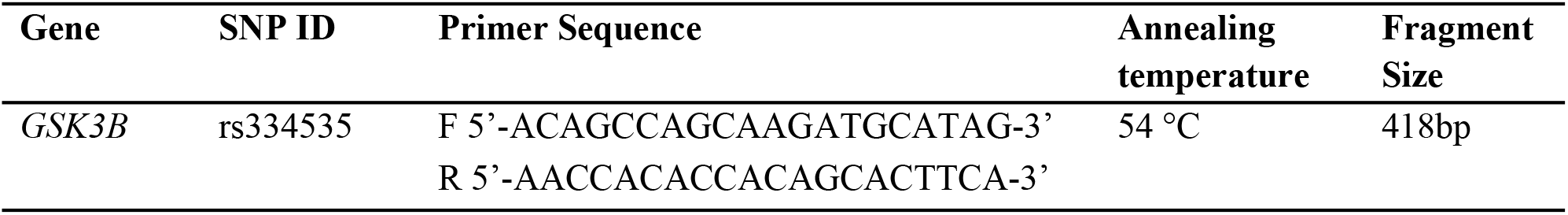
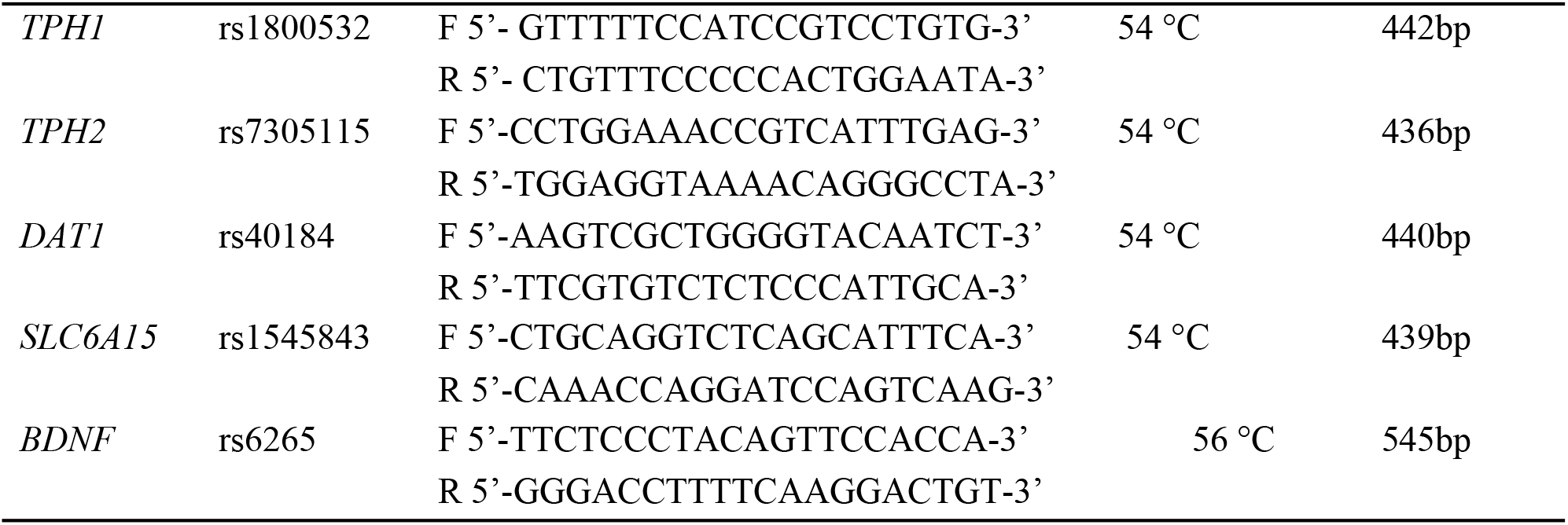
Primers for PCR amplification.

### Statistical analysis

The Hardy Weinberg equilibrium was assayed using the Chi square test from the Hardy– Weinberg equilibrium for association in the entire sample, and both in depressed individuals and controls. The chi-squared test was used to compare genotypes and allele frequencies between groups. The significance level for all statistical tests was 0.05.

## Results

The genotype frequency was found to be in Hardy–Weinberg equilibrium in the entire study samples in both cases and controls (*P*>0.05 in all cases). Table 4 shows the data for genotype and allele frequencies. As shown in Table 4, there were no significant differences between the case groups and the control group in the allele or genotype frequencies of each polymorphism. To study the association between MDD and polymorphisms of the GSK3B gene, we found genotype (χ2 = 1.16; P = 0.560) and allele (χ2 = 0.78; P = 0.376) frequencies of SNP, rs334535 showed no significant difference between patients and control groups. Based on the calculated Chi square and P value for the genotype frequencies of rs1800532 of TPH1 gene, showed no significant differences amongst the MDD patients and controls group, genotype (χ2 = 10.6; *P =* 0.005) allele (χ2 = 6.40; *P =* 0.011). The genotype distributions of rs7305115 of TPH2 gene, (χ2= 16.6; *P*= 0.000) and allele frequencies (χ2= 8.46; *P*= 0.004) were not significantly different between MDD and normal controls. Furthermore, there were no significant difference in the genotype distribution between MDD and normal controls (χ2= 13.6; *P*= 0.001) or allele frequency (χ2= 7.05; *P*= 0.008) of rs40184 of DAT1 gene. There was no difference between the MDD and without depression in genotype distribution (χ2=13.5; *P*= 0.001) or allele frequency (χ2= 7.23; *P*= 0.007) of the rs1545843 polymorphism of SLC6A15 gene. There was also no significant difference in the genotype distribution (χ2= 19.3; *P*= 6.554) or allele frequency (χ2=40.13; *P*= 2.377) of the rs6265 of BDNF gene between MDD and controls. The frequency of the A allele of all SNPs was higher in patients than in controls.

**Table 4.** Genotypic Frequency of the Polymorphisms in Patients of MDD and Control Groups.

| Genotype<br>Frequencies | SNP,s | Cases | Controls | $\chi^2$ | Df | P |
| --- | --- | --- | --- | --- | --- | --- |

|  |  | *N* | % | *N* | % |  |  |  |
| --- | --- | --- | --- | --- | --- | --- | --- | --- |
| *GSK3B* | rs334535 |  |  |  |  |  |  |  |
|  | AA | 21 | 24.7 | 17 | 42.5 | 1.16 | 2 | 0.560 |
|  | AG | 46 | 54.1 | 10 | 25.0 |  |  |  |
|  | GG | 18 | 21.2 | 13 | 32.5 |  |  |  |
| *TPH1* | rs1800532 |  |  |  |  |  |  |  |
|  | AA | 25 | 50.0 | 11 | 36.6 | 10.6 | 2 | 0.005 |
|  | AC | 15 | 30.0 | 9 | 30.0 |  |  |  |
|  | CC | 10 | 20.0 | 10 | 33.3 |  |  |  |
| *TPH2* | rs7305115 |  |  |  |  |  |  |  |
|  | AA | 32 | 42.6 | 23 | 46.0 | 16.6 | 2 | 0.000 |
|  | AG | 26 | 34.6 | 12 | 24.0 |  |  |  |
|  | GG | 17 | 22.6 | 15 | 30.0 |  |  |  |
| *DAT1* | rs40184 |  |  |  |  |  |  |  |
|  | AA | 25 | 50.0 | 13 | 40.6 | 13.6 | 2 | 0.001 |
|  | AG | 12 | 24.0 | 11 | 34.3 |  |  |  |
|  | GG | 13 | 26.0 | 8 | 25.0 |  |  |  |
| *SLC6A15* | rs1545843 |  |  |  |  |  |  |  |
|  | AA | 22 | 44.0 | 15 | 50.0 | 13.5 | 2 | 0.001 |
|  | AG | 15 | 30.0 | 8 | 26.6 |  |  |  |
|  | GG | 13 | 26.0 | 7 | 23.3 |  |  |  |
| *BDNF* | rs6265 |  |  |  |  |  |  |  |
|  | GG | 53 | 58.8 | 25 | 50.0 | 19.3 | 2 | 6.554 |
|  | GA | 22 | 24.4 | 15 | 30.0 |  |  |  |
|  | AA | 15 | 16.6 | 10 | 20.0 |  |  |  |

**Table 5.**
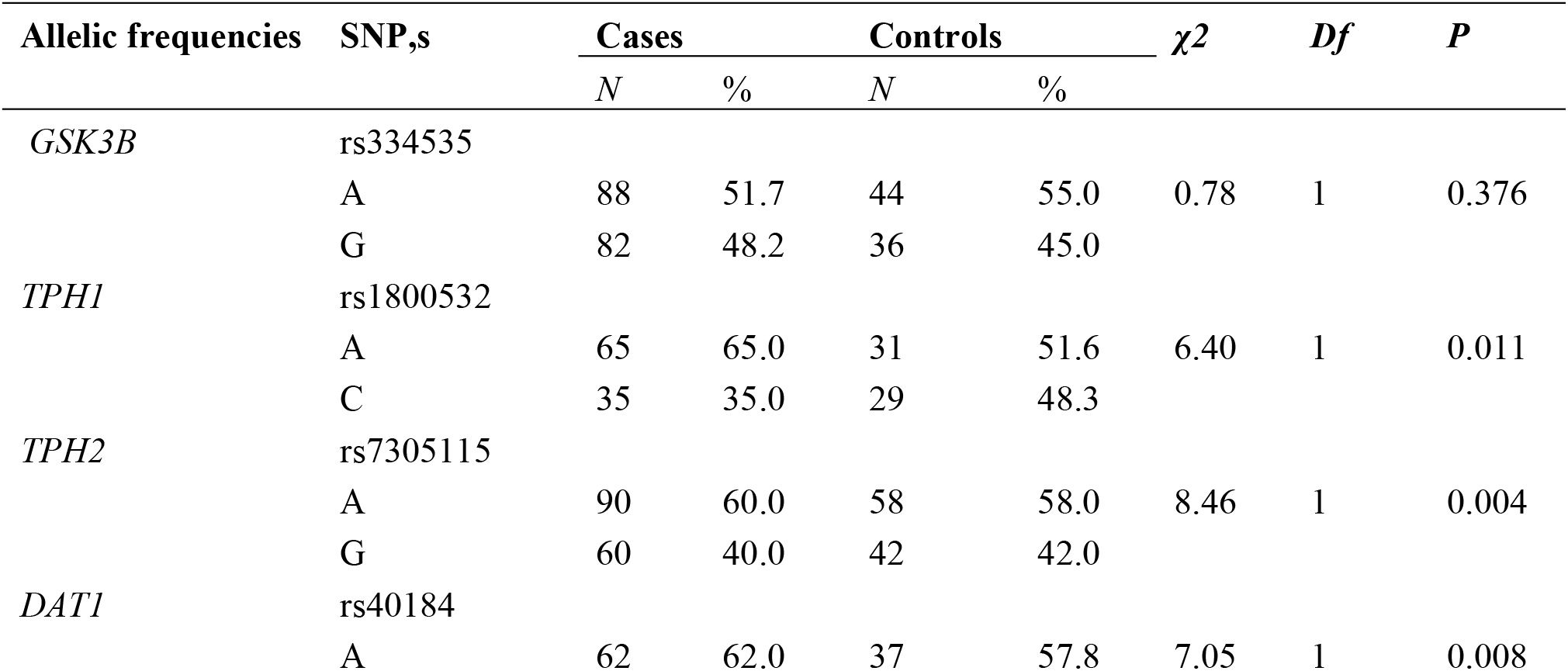
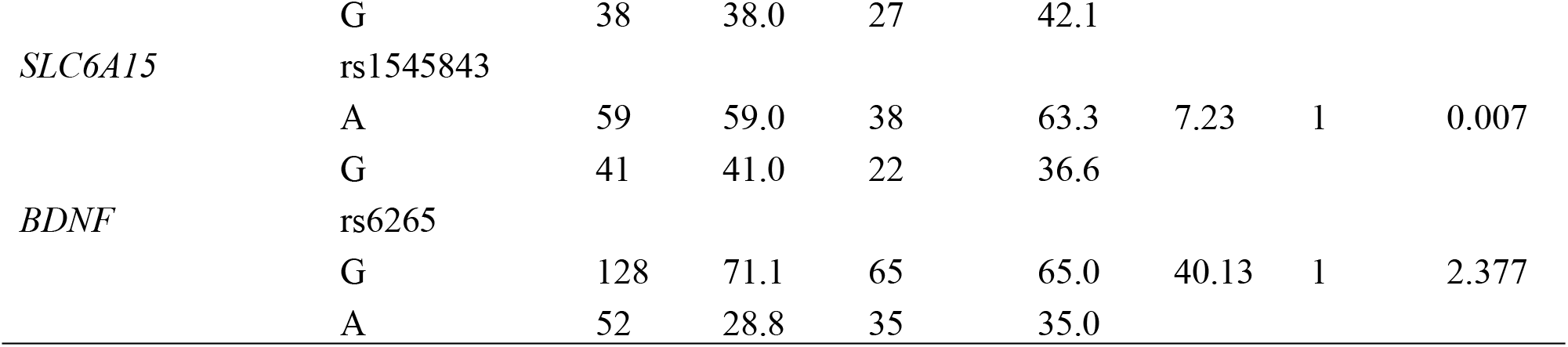
Allelic Frequency of the Polymorphisms in Patients of MDD and Control Groups.

## Discussion

The advancement of genetic studies and knowledge the specific genes, physiological interference responsible in the onset of MDD very limited and is still unclear. Most of the candidate studies only focusing on gene involvement in neurotransmitters circuits or stress reactions which only provided suggestive results although not definitive results. One of the explanations may be phylogenetic nature of the disorder and the limited impact of hypothetically high number of involved loci. In addition, MDD tends to be less homogeneous across population then other psychiatric disorder as bipolar disorder and schizophrenia [12]. To robust the association of such association’s reports independent studies on samples is necessary in current study samples of general population aimed to replicate as previously reported association between candidate gene and MDD of North-Western population (Khyber Pakhtunkhwa of Pakistan). Best to our knowledge it is first ever study examining the association of single SNPs in six genes candidate (*GSK-3B, TPH1, TPH2, DAT1, SLC6A15, and BDNF*) in North-Western population. 400 MDD patients samples were compared with 232 control samples in KP; all SNP’s were calculated through HWE. The GSK-3β gene maps to chromosome 13q13.319 and is located in the same region as the dopamine receptor D3 gene, D3 receptors were localized to limbic areas and may related to cognitive, emotional and endocrine functions [13]. Several lines of evidence show GSK-3B is a good candidate gene for MDD susceptibility. As to date not all studies show that GSK-3B itself is responsible for MDD but with unique clinical characteristics [14]. In current study we investigated differences in GSK-3β gene polymorphisms among patients with MDD, according to individual genetic differences. No difference in the rs334535 polymorphism was found between MDD and normal controls. Our current findings are similar to that of a previous report which was conducted to know genetic interaction with MDD but no significant interaction was founded [15], which strongly agreed with our current study. In contrast a study suggests that interaction between genes can also play an important role in onset of MDD [16], a study conducted in China on Han Chinese population the GSK-3B gene SNP rs334535 was founded to be associated with MDD [17]. Thus, there is a need to investigate other GSK-3β gene polymorphisms and the links between genetic polymorphisms and clinical features of psychiatric diseases.

Tryptophan hydroxylase (TPH) is responsible for serotonin and neurotransmitter synthesis and coded ahead by TPH1 and TPH2 both isoforms have same 70% sequence identity but the pattern of expression is different and both works on rate-limiting and the biosynthesis of serotonin. TPH1 isoform is predominant in periphery and the TPH2 in central nervous system [18]. Polymorphism of TPH1 gene is subjected for different kind of researches. Most known variation of this gene is A218C SNP (rs1800532) of intron 7 [19]. Different studies also showed that while expression of TPH1 gene the allele A of A218C play an important role [20]. In present study genotyping of TPH1 polymorphism in MDD patients and controls were not in agreement with the Hardy-Weinberg equilibrium (HWE). A study reported that genotype CC showed association to suicide attempts in individuals aged more than 65 years [21]. Our current study didn’t show any kind of association related to suicide and the association was not age related polymorphism of A218C TPH1 gene know to be linked with different disorder like Borderline personality disorder (BPD) in unites states [19], suicidal behavior and anger-related personality behaviors [22], schizophrenia [23], bipolar disorder in Taiwan [24], Caucasian population [25] and depression in Finland [26]. Different association studies were conducted on TPH1 A218C polymorphism in Germany and China and were failed to find any association [27], there is lot of contradiction founded in this area.

TPH2 localized at (12q21.1) discovered after TPH1 play important role in neural tissue that’s why called brain specific isoform [28]. Compare to TPH1 it express more in brain that’s why it is suggested it has more significant role in brain region. Different studies reported variation of this genes leads to suicidal attempts, MDD and Seasonal Disorder [29,30], autism [31], and schizophrenia [32]. Finding of our current study TPH2 gene polymorphism (rs7305115) did not show any association with MDD. While comparing to previous studies the current finding is agreed with lots of studies. Association of allelic and genotypic frequencies in TPH2 in MDD patients [29,33], a study related to gender base do not showed a significant result in TPH2 gene effect on susceptibility of MDD [34,35], another study target single locus and haplotype analysis didn’t show any association while targeting SNP rs7305115 in Japanese population [36].

Though the cellular level of DAT1 gene and this SNP rs40140 level have not demonstrated yet previous studies linked the polymorphism of unknown function to some psychiatric diseases [37]. So we conduct a study on genotypic distribution, allelic frequencies allele carrier frequencies on rs40184 same results was found no significant differences founded between groups case and control groups as well as in allelic frequencies, genotypic distribution and allele carrier frequencies in this particular SNP. Similar study carried out and founded no significant differences analyzing the same way but they observed more carriers C allele (CC+TT genotypes) in control when compared to patients so it suggest tendency toward association between genotype and TT homozygous this can be increased risk of MDD [38]. Previously another study reported that Russian population those reported with maternal rejection also had TT genotype of SNP rs40184 [39], founded to be settled with our study.

SLC6A15 gene is founded a novel candidate gene responsible for MDD still it is unknown how SLC6A15 genes alter brain functional activities of MDD patients. The SLC6A15 coding for a sodium dependent branched amino acid transporter is the nearest annotated gene with a distance of approximately 287kb to the region of association [40]. In the present study we investigate the impact of polymorphism of SLC615 gene rs1545843 found no significant association between case and control samples. Recently SLC6A15 has been found to be MDD associated particularly in a genome wide association analysis, SNP rs1545843 documented with significant result [40]. Another study found an effect of rs1545843 in MDD patients on adrenocorticotropic hormone (ACTH) and cortisol levels and documented an association in these patients between his polymorphism and cognitive functions such as memory and sustained attention [41]. Although a recent GWAS investigated the proposed role of this genes in the stress susceptibility documented in animal model studies and potential associations with a predisposition to MDD [40]. The exact neurobiological role of this genetic polymorphism in the pathophysiologic phase of MDD and association of structural changes in specific brain regions in not clearly explained.

The human BDNF gene was mapped to 11p13 chromosomes in a coding exon of the BDNF gene at position 66 (Val66Met, rs6265) a common single nucleotide polymorphism (SNP) consisting of a missense change (G196A) causing a non-conservative amino acid conversion (valine to methionine) was established. The genetic relationship between BDNF and MDD has been examined in a substantial amount of literature nevertheless the BDNF polymorphism is associated with an increased risk of depression reported in many studies [42], while some studies failed to find the association [43]. In our current study using single loci analysis BDNF rs6265 was not found to be correlated with occurrence of MDD studied in KP population Pakistan. Several researchers reported the interaction between the BDNF gene and MDD stating that both pathophysiology and MDD diagnosis are extensively heterogeneous and thus this character makes them difficult to detect a susceptible gene for MDD [27,44]. There is several interpretation and discrepancies in addition MDD is complex disorder that is believed to be caused by effects of various genetic factors in the pathophysiology of the disease and gene-gen interactions are likely to contribute.

## Conclusion

Our primary goal of finding interacting genetic risk loci for major depressive disorder susceptibility has not been successful. Despite the lack of significant results, we recommend that the current study still contain interesting results. Firstly, with restriction to the six candidate genes our results further support the hypothesis that MDD has multifactorial background with an exceptionally evolved network of genetic interactions contributing small amount to the overall- risk of developing MDD. The field of psychiatric genetics is generally had been disappointing, because the initial hope of finding common genetic variants that play an important role in the pathogenesis of mental illness has not been successful. In most psychiatric illnesses, the phenotype seems too complex, with the patient cohorts too small and no findings have been consistently replicated. This is also the case for MDDs. In addition, the phenotypic effects of genetic variants identified to date are weak. When comparing the impact of genetic variation on disease susceptibility and the impact of lifestyle and environmental factors, the situation is more complex, and the impact of lifestyle and environmental factors may be significant. Despite these obstacles, the field of psychogenetic is still developing rapidly, and some new technological advances have been made (such as whole genome sequencing, GWAS). It is important to remember that genetic information will only provide additional information about one aspect of a psychiatric patient’s complex and personal history. It is the sum of internal and external factors that affect and influence mental health and well-being. Therefore, more research is needed to better cover these six candidate genes and explore other variants that may affect gene expression. GWAS studies and large scale analysis or additional work in different populations and appropriate analytical analysis are still needed to determine conclusively the etiology of MDD and other mental illnesses. Identifying these interactions will help to prioritize biological processes that require further study to better understand the etiology of MDD.

## Acknowledgments

We are sincerely thanks to the patients, their families, and the healthy volunteers for their participation, and all the medical staff involved in specimen collecting, as well as those who contributed to this work in Human Genetics Lab, Hazara University Garden Campus, Mansehra 21300 Pakistan.

## Data Availability

All data generated or analyzed during this study are included in this article (and are available from the corresponding author on reasonable request).

## Author contributions

1. Naqash Alam: Help in lab experimentation and acquisition of data analysis
2. Sadiq Ali: Help in lab experimentation and acquisition of data analysis
3. Nazia Akbar*: Execute basic concept and design research work Muhammad
4. Ilyas: Help in study design and proof reading of this article
5. Habib Ahmad: Provide administrative and technical support
6. Arooj Mustafa: Help in lab experimentation and sample collection
7. Shehzada Khurram: Help in lab experimentation and sample collection
8. Zeeshan Sajid: Help in lab experimentation and sample co
9. llection Najeeb Ullah: Help in lab experimentation and sample collection
10. Shumaila Qayyum: Help in lab experimentation and sample collection
11. Tariq Rahim: Help in lab experimentation and sample collection
12. Mian Syed Usman: Help in lab experimentation and sample collection
13. Nawad Ali M: Help in lab experimentation and sample collection
14. Imad Khan: Help in lab experimentation and sample collection
15. Khola Pervez: Help in lab experimentation and sample collection
16. BiBi Sumaira: Help in lab experimentation and sample collection
17. Nasir Ali: Help in lab experimentation and proof reading of this article
18. Nighat Sultana: Critical revision of the manuscript
19. Adeel Yunus Tanoli: Critical revision of the manuscript
20. Madiha Islam: Critical revision of the manuscript

